# Global biogeography since Pangaea

**DOI:** 10.1101/146142

**Authors:** Sarah R. N. McIntyre, Charles H. Lineweaver, Colin P. Groves, Aditya Chopra

## Abstract

The breakup of the supercontinent Pangaea over the past ~ 180 million years has left its imprint on the global distribution of species and resulted in vicariance-driven allopatric speciation. Here we test the idea that the molecular clock dates for the divergences of species whose geographic ranges were divided, should agree with the palaeomagnetic dates for the continental separations. Our analysis of recently available phylogenetic divergence dates of 42 pairs of vertebrate taxa, selected for their reduced ability to have undergone dispersal-driven speciation, demonstrates that the divergence dates in phylogenetic trees of continent-bound terrestrial and freshwater vertebrates are consistent with the palaeomagnetic dates of continental separation.

---

When a landmass breaks up due to continental drift, the geographical ranges of thousands of species are simultaneously divided^1^. Vicariance-driven allopatric speciation is followed by radiation on each side of the breakup. Subsequently, over millions of years, pairs of geographically separated taxa emerge from the original species. Each pair of these sister taxa will have a divergence time from a common ancestor that should correspond to the time of the continental breakup. The recent explosion in the number of species with sequenced genomes includes many species that belong to the geographically separated sister taxa. These sequences permit construction of well-calibrated phylogenetic trees and molecular-clock-based estimates of the divergence times of those taxa^2,3^. Our goal is to verify that divergence dates based on molecular clocks correlate with palaeomagnetic dates for the continental separations, and to help quantify the extent of their agreement.

Previous palaeobiogeographic research has been either taxon specific (e.g. Platnick’s spiders,^4^ Mittermeier’s birds,^5^ and Zhang et al.’s frogs;^6^), or continent specific (e.g. Tabuce et al.’s Afrotheria,^7^ Gibb et al.’s Xenarthra,^8^ and Hu et al.’s Laurasiatheria^9^). Here, we analyse the global chronology of biogeographic divergences from published phylogenies^2,3^ of 42 vertebrate taxa pairs and compare them with palaeomagnetic dates for the breakup of Pangaea.

Pangaea began to separate ~180 million years ago (Fig. 1) into a southern continent, Gondwana (that contained what is now South America, Africa, India, Madagascar, Australia, New Zealand and Antarctica), and a northern continent, Laurasia (that contained what is now North America, Europe and Asia, minus the Indian subcontinent). Subsequently, Gondwana and Laurasia each split to produce the present biogeography. Thus, the morphological similarities of pairs of taxa from South America and Africa, are not the result of the dispersal of several hundred species across the Atlantic by rafting, but instead result from both faunas being descendants of the same ancestral biota that split ~100 million years ago.

**Figure 1.**
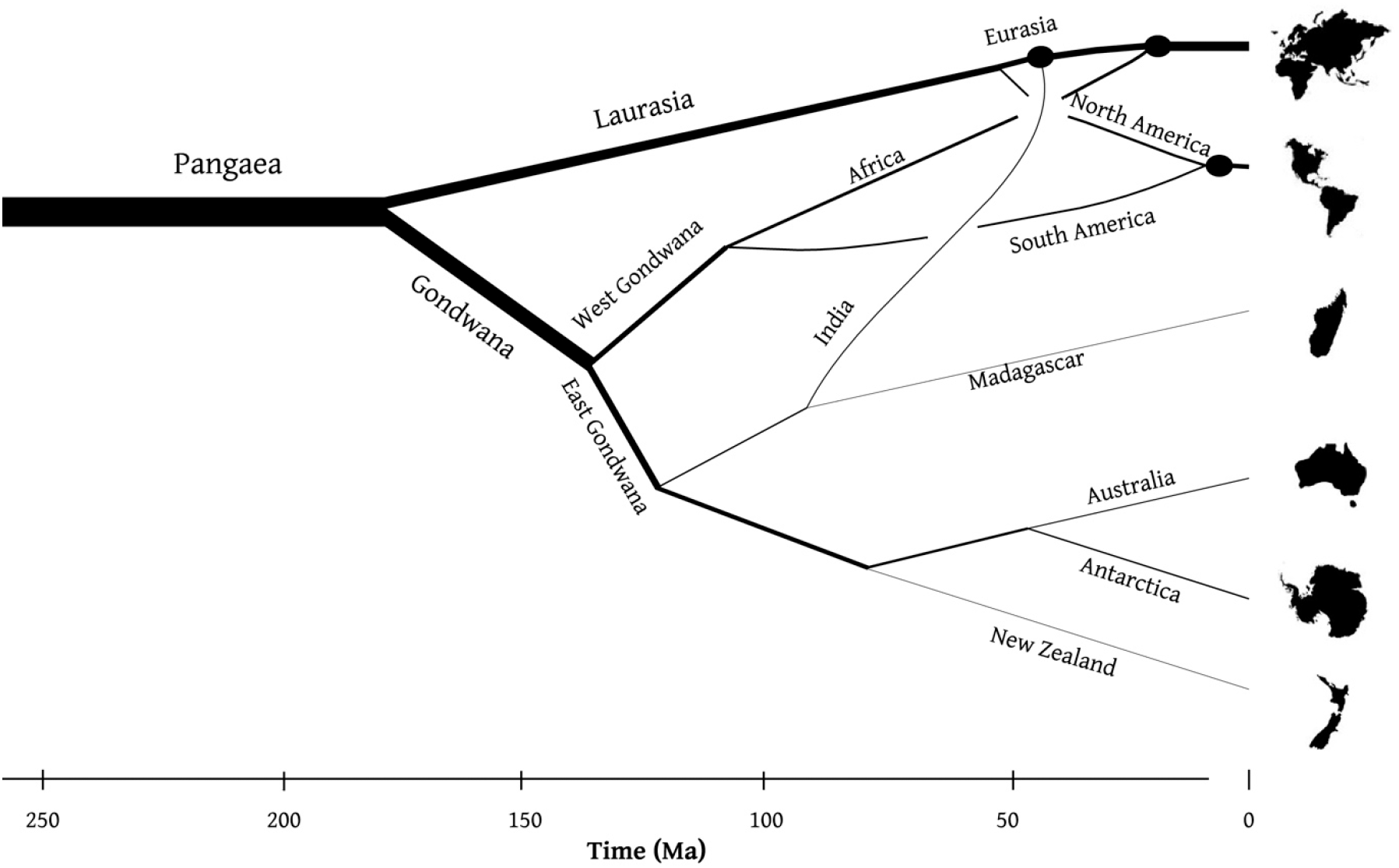
Breakup of the supercontinent Pangaea over the past ~180 million years. Geological and biological chronologies provide two independent ways to date 8 continental divergences and 3 continental convergences. Continental separations are not instantaneous and occur over tens of millions of years (Supp. Table 1, column 7). Such durations are comparable with the amount of correlation between the palaeomagnetic and phylogenetic divergence dates and set a lower limit to the expected level of agreement. Circles indicate continental collisions. Line thickness is a rough proxy for landmass area.

The plate tectonic history of Pangaea has been well-documented using palaeomagnetic dating. The chronology of the breakup of Pangaea presented here (Fig. 2a) is based on the weighted average of four palaeomagnetic data sources for each of 8 continental separations and 3 continental collisions (Supp. Table 1).

**Figure 2.**
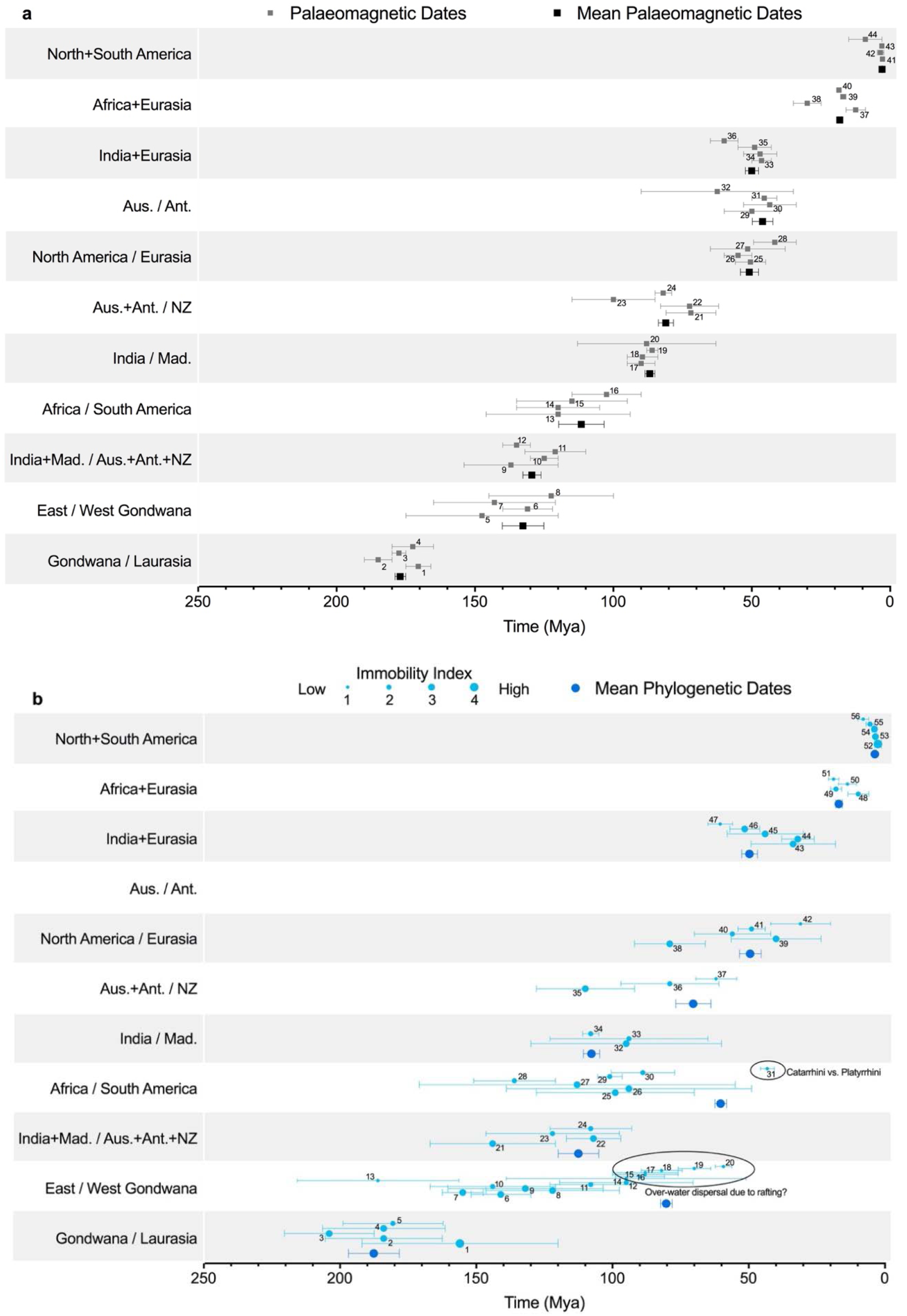
**a.** Palaeomagnetic dates for continental divergences and collisions. Numerical labels (1 - 44) refer to references (Supp. Table 1). In **b.** we plot molecular clock divergence dates of allopatric speciation associated with continental divergences (1 - 42) and biotic interchange dates associated with continental collisions (43 - 56) (Supp. Table 2). No phylogenetic data from the Australia/Antarctica divergence is applicable; thus, this divergence is represented by only palaeomagnetic data. Point sizes of phylogenetic data indicate the immobility index for that pair of taxa (Table 1 and Supp. Table 2).

Thousands of species have had their geographical ranges divided by continental movements. Most of these species went extinct either on one or both sides of the division. Therefore, extant endemic sister taxa, on separated continents, were required to analyse the vicariance phenomenon. Some species could have diverged before continental breakup; however, with small number statistics, these are difficult to identify. Fossil-based attempts at a global biogeographic analysis have been compromised due to the sparseness of the fossil record and the fact that fossils (unlike molecular clocks) usually provide only minimum age constraints on continental divergence times^10^. The timetree.org database^2,3^ has assembled a comprehensive sample of phylogenetic molecular clock divergence dates from current literature, which have been used in our analysis (Supp. Table 2).

Molecular divergence dates from 42 pairs of vertebrate sister taxa (labelled 1 - 42), derived from 74 separate taxa on the isolated continents, are shown in Fig. 2. The remaining 3 continental collisions (India+Eurasia, Africa+Eurasia, North+South America) were biologically evaluated using the biotic interchange data from 14 vertebrate taxa (labelled 43 - 56). While the most common evidence for the collision events was fossil data, endemic species dispersal to the new continent and their subsequent diversification (traced phylogenetically) was also indicative of a migration event.

It was important that phylogenetic divergences in our analysis were not due to dispersal (e.g., by rafting or island hopping), but rather, due to ancient vicariant events related to plate tectonic movements. For example, a strong case exists for multiple instances of rafting between Africa and Madagascar.^11–13^ Several Late Oligocene to Mid Eocene (~26 - 50 Mya) terrestrial mammalian groups in Madagascar are closely related to those found in Africa.^14^ Molecular clock data indicates that the tenrecs would have begun diversification at 20 - 32 Mya, the nesomyines (Malagasy rodents) at 18 - 30 Ma, and the Malagasy carnivores at 14 - 25 Mya.^15^ One proposal regarding the origin of these mammals on Madagascar suggests the brief existence of a land-bridge, along the Davie Ridge fault line, during this period.^15, 16^

The less mobile the species, the more likely its divergence into two species resulted from continental separation. To minimise the effect of dispersal between continents, taxa were selected for their reduced ability to have undergone dispersal-driven speciation. Therefore, we have selected continent-bound terrestrial and freshwater vertebrates. Additionally, we have created an immobility index to help quantify the extent to which each of the vertebrate taxa were continent bound. The mobility of the taxa depends on many intrinsic (e.g. body condition) and extrinsic (e.g. climate) factors.^17^ The four immobility factors described in Table 1 were assessed for each of 56 vertebrate taxa (Supp. Table 2).

**Table 1.** Immobility factors

| Immobility factors | Reasoning | Example |
| --- | --- | --- |
| Dispersal restraints | The species would need to endure a sea voyage including exposure to saltwater and fluctuating temperatures. If the species is salt or temperature sensitive, it might not survive long enough to disperse across and populate the new continent. | Freshwater fish and selective frogs |
| Dietary requirements | For a species to survive crossing a continental divide it would need to be flexible enough with its diet to find satisfactory food sources on the sea voyage and upon arrival in the new ecosystem. Species in a commensal or symbiotic relationship would have difficulty adapting without their partners. | Specific fish, geckos and caecilians |
| Reproduction limitations | Some animals reproduce rapidly and produce many offspring (*r*-selection). With only one pregnant female, these animals could rapidly colonise a new continent. Others do not reach sexual maturity for many years after birth and then produce few offspring (*K*-selection) and would be less likely to colonise a new continent. | Primates and mammals |
| Mating limitations | Seasonal breeders successfully mate only during certain times of the year. These mating periods result in births at times optimal for survival of the young, taking into account factors such as ambient temperature, food and water availability, and changes in predation behaviours of other species. | Selective mammals, fish and frogs |

Continent-bound, immobile species, were assigned a high immobility index of I = 4. More mobile species (that might be able to cross an ocean) were assigned an immobility index of I = 1. We estimated the immobility index for each pair of taxa (Supp. Table 2) without reference to its divergence date. Then, in Fig. 2b, we plotted divergence dates and used point size to indicate the immobility index.

Inclusion of mobile species able to navigate across broad ocean channels and survive in diverse environments would systematically shift the biology-based divergence to more recent times.

This tendency can be seen in the points labelled 15 through 20 (Fig. 2b), in the East/West Gondwana divergence (inside the oval labelled “Over-water dispersal due to rafting?”) for which I=1. Another outlier is the Catarrhini/Platyrrhini taxa pair (31) associated with the Africa/South America divergence. These are highly mobile primates that were assigned a low immobility index I = 1. Separation of these 2 taxa could be ascribed to over-water dispersal millions of years after continental split. On the opposite side of the immobility spectrum the African torrent frog vs neotropical true frog (30) was assigned a high immobility index because, in general, frogs are intolerant of salt water. The taxa pair: Gymnopis (American caecilians) vs. Ichthyophis (Asian caecilians) in the Gondwana / Laurasia divergence, is an example of the most immobile taxa. Their high immobility is due to their burrowing lifestyle and intolerance of salt water in which, like most amphibians, they suffer desiccation. For continental collisions, as continents moved closer together, more mobile species could have dispersed before the collision occurred. Thus, I = 1 points in collisions should have older dates. A slight tendency in this direction is seen for the I = 1 points for the 3 collisions at the top of Fig. 2b.

We have combined the data in Fig. 2 to produce Fig. 3 with the following modifications. All I = 1 points were excluded, and the immobility index was used to modify variance of the remaining biology points such that when computing the weighted average, mobile taxa were down-weighted and immobile taxa had more weight (Eq. 1).

**Figure 3.**
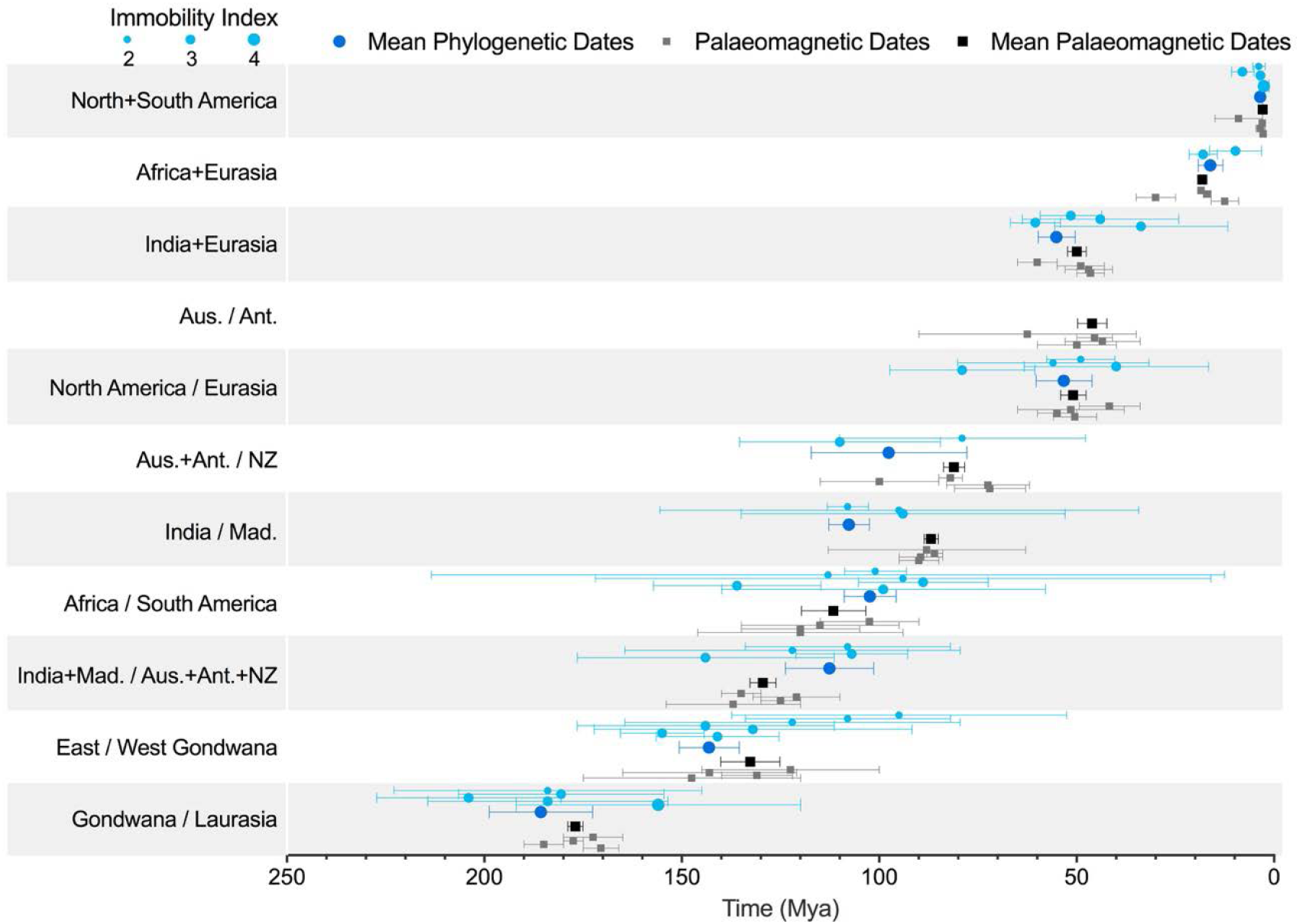
Phylogenetic and palaeomagnetic data modified from Fig. 2. The most mobile taxa with immobility index I = 1 are excluded. We used the immobility index (through Eq. 1) to modify variances for computing phylogenetic mean values.

In Fig. 4 time averages from palaeomagnetism (x-axis, black horizontal error bars) and phylogeny (y-axis, blue vertical error bars) are highly correlated (0.98 correlation coefficient). For most points, uncertainties for the palaeomagnetic values are smaller than analogous phylogenetic uncertainties. However, notable exceptions include the approximately equal length of both vertical and horizontal error bars for East / West Gondwana and Africa / South America. Increases in the number of species with full genomes and improvements to molecular clock calibrations should reduce uncertainties on the phylogeny-based divergence times. A more targeted study of rafting cases and potential land-bridges should be conducted to pinpoint whether the phylogenetic divergences are due to vicariance or dispersal. Additionally, there is an ongoing polemic over whether New Zealand was above sea-level for the entirety of its isolation, or was submerged, and only populated by dispersal upon reemergence. Uncertainties in the palaeomagnetic dates are also decreasing^18^.

**Figure 4.**
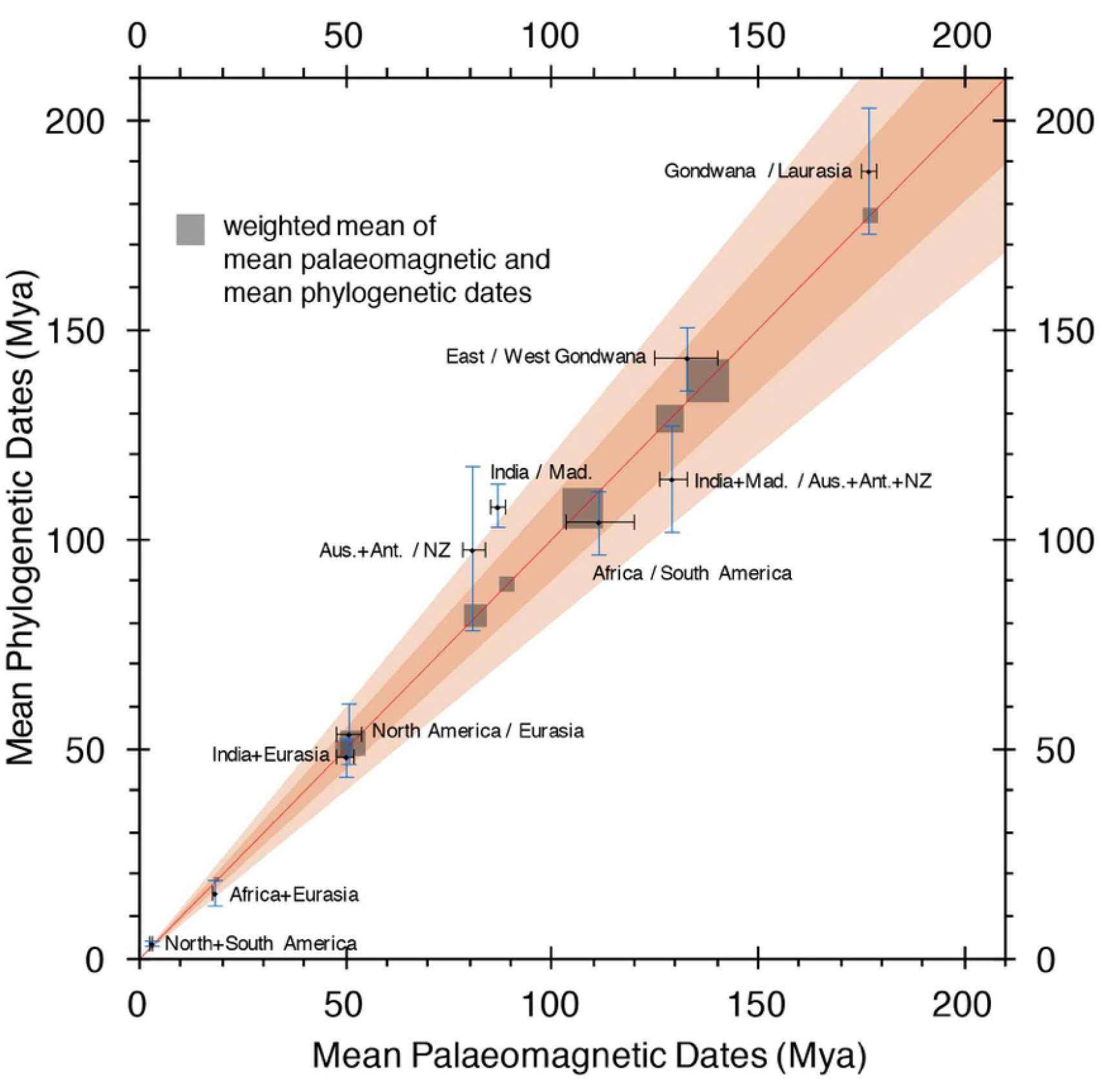
Comparison of the mean palaeomagnetic dates with the mean phylogenetic dates. Weighted means of the two independent estimates are represented by grey squares along the diagonal (Supp. Table 3). With perfect agreement, all points would lie on the red line with slope=1. Slopes of the dark and light orange wedges are 1 ± 0.1 and 1 ± 0.2 respectively. Error bars in the vertical and horizontal directions for East / West Gondwana and Africa / South America have approximately equal length. Thus, phylogenetic data are becoming of equal importance in dating continental divergences.

With continually improving techniques for estimating both palaeomagnetic continental breakup dates and molecular-clock-based phylogenetic dates a more precise and accurate comparison can be made, not only for Pangaea but potentially for the breakup of earlier supercontinents, e.g. Pannotia (~ 545 - 600 Mya) and Rodinia (~ 1.1 - 0.75 Gya).

## Methods

### Immobility Index

Our immobility index assigns values between 1 and 4 - with 1 assigned to the most mobile taxa and 4 to the least mobile. When computing weighted averages in Fig. 3, we exclude pairs of taxa with immobility 1, and assign more weight to pairs with the highest immobility, since they have divergence dates that are less likely to result from dispersal. Specifically, we increased the reported variance of a phylogenetic divergence date 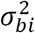 depending on the immobility index as follows:

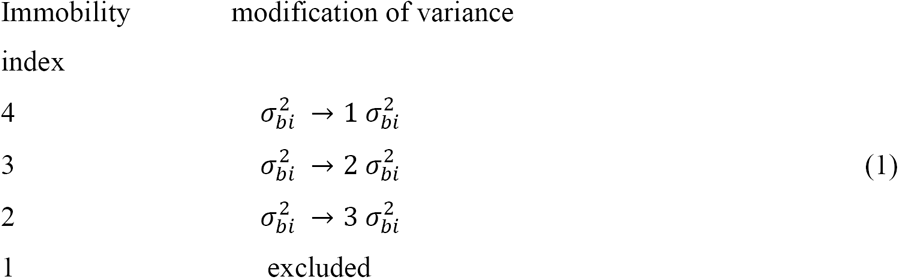

With these modifications to 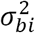 we computed inverse variance-weighted means in the same way as for the geological data. Let *g*_*i*_ and b_*i*_ be the geology and biology inverse-variance-weighted average dates for the *i*th continental break-up (or collision). Let 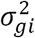 and 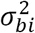 be their variances respectively (after the 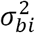 have been modified as described in Eq. 1). Let *t*_*i*_ be the weighted average of *b*_*i*_ and *g*_*i*_:

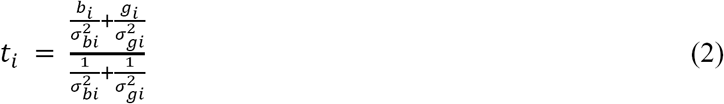

### Chi-Squared Test

We used χ^2^ per degree of freedom (χ^2^/dof) tests to provide some justification for our reported uncertainties. For example, the palaeomagnetic literature reports ranges that we take as “1 sigma” ranges or 68% confidence intervals. For each of the 11 nodes in Supp. Table 1, we compute χ^2^/dοf The average of these 11 values is ~ 1. Similarly, χ^2^/dof values for the 10 nodes in Supp. Table 2 (after removal of the I=1 data) suggest that uncertainties reported on the TTOL (Time Tree Of Life) values in timetree.org can be plausibly taken as 68% confidence intervals. To determine whether *b*_*i*_ and *g*_*i*_ are consistent with *t*_*i*_ we compute χ^2^ (Eq. 3) and verify that χ^2^/dof ~ 1.

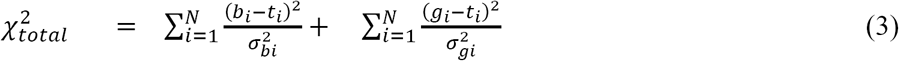

where N = 10. See the last two columns of Supp. Table 3.

## Acknowledgements

We thank Stephen Blair Hedges, Andrew Roberts and David Heslop for helpful discussions and correspondence.

## Author Contributions

S.R.N.M and C.H.L did the analysis. C.P.G and S.R.N.M were responsible for the immobility index. All authors contributed to the writing of the manuscript.

## Author Information

Correspondence and requests for materials should be addressed to

